# Interactions between time on diet, antibiotic treatment, and liver abscess development on the fecal microbiome of beef cattle

**DOI:** 10.1101/2022.09.16.508319

**Authors:** Germán Plata, Nielson T. Baxter, Troy B. Hawkins, Lucas Huntimer, Akshitha Nagireddy, Dwi Susanti, James B. Reinbold

## Abstract

Liver abscesses caused by polymicrobial infections of the liver are a widespread problem in feedlot cattle production. There are currently no effective methods for the early detection of liver abscesses or to predict antibiotic efficacy. Although gene expression and microbiome differences have been reported in the rumen of abscessed and normal animals, liver abscess biomarkers using less invasive tools can facilitate managing of the disease in the field. Here we report the results of two studies measuring the fecal microbiome composition of steers that did or did not develop liver abscesses, with or without antibiotic treatment, along a 7-month feeding period on a high-concentrate diet. Our results indicate a limited impact of liver abscesses or tylosin on fecal microbiome composition, with time on diet explaining most variance in the fecal microbiome. Interestingly, in both studies, antibiotic treatment led to larger differences in both the composition and variance of the fecal microbiomes between abscessed and normal animals compared to controls. These differences were limited to specific sampling times in each of the two studies. Although multiple amplicon sequence variants with differential abundances according to liver abscess state were identified, there was no overlap between the two studies. Our results suggests that early fecal biomarkers of liver abscess susceptibility might be developed, especially for animals receiving preventative antibiotics, but the fecal abundance of individual microorganisms may not be a robust predictor across different sampling times or diet regimes.

## Introduction

High-energy diets fed to cattle in the feedlot are an important driver of performance but may also negatively affect animal health if not properly managed (1, 2). Liver abscesses are pus-filled pockets in the liver of cattle that form due to bacterial infection and are often a direct result of feeding practices (3), resulting in liver condemnation, performance losses, and reduced carcass weights (4, 5). It is hypothesized that the rapid fermentation of grain-rich, low roughage finishing diets contributes to acidosis-driven lesions in the bovine gastrointestinal (GI) tract, enabling the translocation of members of the GI microbiome such as the opportunistic pathogen *Fusobacterium necrophorum* into the bloodstream and liver of animals that develop liver abscesses (3, 6).

While feeding high levels of roughage during the growing and finisher periods can reduce the prevalence of liver abscesses (7), the most common method for liver abscess control has been the continuous feeding of antimicrobial compounds (6, 8). For instance, tylosin phosphate has been shown to reduce the load of *F. necrophorum* in the rumen (9), and lower the prevalence of liver abscesses by more than 50% compared to untreated animals (10-12). Nevertheless, how tylosin or other antibiotics may prevent liver abscesses, or why protection levels vary between animals, are not fully understood. As global initiatives to minimize the use of antimicrobials for livestock production are underway (13, 14), a greater understanding of when and why these drugs are effective would benefit efforts to enhance their application.

Abbas et al. recently demonstrated differences in gene transcript levels in the rumen epithelium, as well rumen-wall associated bacterial communities, between animals with or without liver abscesses (15). Macdonald et al. showed associations between liver abscesses and blood analytes collected up to 56 days prior to slaughter (16). While these results suggest it may be possible to develop diagnostic tests for liver abscess prior to slaughter based on rumen or blood biomarkers, an association between fecal biomarkers and the occurrence of liver abscesses would further facilitate diagnosis and management of the disease given an easier access to fecal compared to rumen or blood samples. Weinroth et al. looked at the fecal and soil microbiomes of cattle fed with or without tylosin at the level of pens, finding that both types of microbial communities could predict 75% of the variance in abscess prevalence per pen (17). Thus, the possibility exists that fecal microbial communities can be used to diagnose liver abscesses in individual animals.

Here we investigate whether the composition of the fecal microbiome of individual steers at different time points after transitioning to a high-grain diet is associated with the presence of liver abscesses at slaughter. We analyze samples from two different studies in which diets were supplied with two different types of antimicrobials. In the first study, cattle were fed with or without tylosin, a macrolide, whereas in the second study cattle were fed with or without narasin, an ionophore. The longitudinal data collected allowed us to investigate the interaction between antibiotic treatment, time on diet, and liver abscess diagnosis on the cattle fecal microbiome. Our results indicate that early differences in the fecal microbiome of animals with or without liver abscesses at slaughter are more common in antibiotic treated steers compared to controls.

## Materials and methods

### Study 1

The live animal experiment and procedures were approved by the Elanco institutional animal care committee, approval number EIAC-0785.

400 Purebred or crossbred steers of British or Continental breed genetic influence were sourced through standard market channels. Animals were placed in feedlot pens and maintained on grass hay for at least 21 days of acclimation until the start of treatment (day 0). Half the animals were randomized to each of 2 treatment groups in a complete block design. The blocking factors were arrival body weight and location within the test facility. Twenty animals were group-housed in a pen, with two pens per block and 10 pens per treatment group. On day 0, the forage-based diet was replaced by a starter ration, from day 5 to day 16 animals were fed a first intermediate diet (“home calf” or similar), followed by a second intermediate diet (“step up” or similar) from day 17 to day 32. From day 33 until harvest animals were fed a finishing diet (“finish” or similar). From day 0, animals in the first treatment group were fed 0 mg of tylosin/head/day (referred to as the S1-control treatment), while animals in the second treatment group received 80 to 90 mg of tylosin/head/day (S1-tylosin treatment). Animals were tracked via ear tags labelled with a numeric individual and group pen identifier.

The arrival weight of the steers was between 250 and 387 kg. Animals appearing visually unthrifty, ill, and/or injured were excluded from the study, as well as animals with ultrasonographic evidence of liver abscess disease or other liver pathology. Animals with a known history of macrolide-or penicillin-class antibiotic use within 60 days prior to purchase as well as a history of vaccination with an anti- *Fusobacterium necrophorum* vaccine were also excluded.

Fecal samples were collected from 43 animals, 21 from S1-control and 22 from S1-tylosin, on days 84, 126, 168 and 220. The 43 sampled animals were selected seeking an even representation of normal and abscessed individuals in both the control and tylosin treatment groups. Animals with liver scars but no clear abscess phenotype were not considered. Preference in sample selection was given to pens with both normal and abscessed animals.

### Study 2

The live animal experiment and procedures were approved by the Elanco institutional animal care committee, approval number EIAC-1017.

The number and type of animals and study design was equivalent to Study 1 above, with differences in the treatments, diet regimes and sampling times. A forage-based diet was provided from animal arrival until prior to the study start on day 0. Animals in the control treatment (S2-control) were fed a starter ration from day 0 to day 10, a first intermediate diet (“home calf” or similar) from day 11 to day 32, a second intermediate diet (“step up” or similar) from day 33 to day 66, and lastly, a finishing diet (“finish” or similar) from day 67 until harvest. Animals in the second treatment (S2-narasin) received the same diets, supplemented with 10g narasin/ton dry matter in the feed.

The arrival weight of the steers was between 233 and 405 kg. Animals were excluded from the study according to similar criteria as study 1.

Fecal samples were collected from the 40 animals (20 from each treatment) assigned to the two pens in the block with the lowest starting weight. These animals were on feed for the longest time prior to harvest. Samples were collected on days -2, 28, 56, 84, 112 and 140 of the study.

### Liver abscess scoring

For both studies, animals were sent for harvest to a commercial slaughter facility and each animal had a liver abscess score assigned. Harvest was conducted according to standard USDA slaughter practices for the facility. At the time of examination, livers were visually scored by a trained observer according to the presence and severity of abscesses or absence of abscesses according to the criteria in table 1. For the analyses, we considered a sample to be from an abscessed animal if necropsy results were in the A-, A or A+ categories.

**Table 1.** Definition of abscess scores assigned to animals upon necropsy.

| Diagnosis |  | Definition |
| --- | --- | --- |
| Normal |  | No abscesses present |
| Abscess | A- | 1 to 2 small abscesses ($\leq 1$ inch in diameter) present |
| | A | 3 to 4 small abscesses ($\leq 1$ inch in diameter) present |
| | A+ | 1 or more large abscesses ($> 1$ inch in diameter) present |

### Sample collection and 16S rRNA sequencing

Fecal samples were collected directly from the rectum via insertion of a lubricated, gloved hand and the removal of a grab sample of feces. One gram of feces was acquired by filling the spoon of a DNA/RNA Shield™ Fecal Collection Tube kit (Catalog No. R110150). The spoon containing the fecal sample was then inserted into the Zymo DNA/RNA Shield solution. The tube was shaken by hand to mix the contents and transferred to a cooler containing cold packs. Gloves were changed between animals to prevent cross-contamination. Samples were transferred from the coolers to long term storage in a -60C freezer on the same day of collection. Samples were maintained frozen until preparation for DNA extraction procedures.

DNA extraction and library preparation were performed using the Shoreline Biome V4 16S DNA Purification and Library Prep Kit (Shoreline Biome, Farmington, CT). DNA was extracted starting from 10 ml of homogenized fecal slurry. The V4 region of the 16S rRNA gene was PCR amplified using the 515F (5’GTGGCCAGCMGCCGCGGTAA) and 806R (5’-GGACTACHVHHHTWTCTAAT) primers following the manufacturer’s protocol. Amplicons were sequenced using 2 × 250 bp paired-end kits on the Illumina MiSeq platform.

### 16S rRNA data analysis

Sequence data from the two studies were generated and processed independently using the following procedure. Paired raw reads were processed with cutadapt (v. 2.5) (18) to remove primer sequences using parameters *-m 10, -e 0*.*15 --discard-untrimmed*; i.e. read pairs that did not contain the primer sequences with a tolerance of mismatches of 15% were discarded. After removal of primer sequences, read pairs were processed using the DADA2 pipeline (v. 1.12.1) (19) to generate a matrix of read counts per sample at the level of amplicon sequence variants (ASVs). Reads were truncated to a maximum length of 250 nucleotides and filtered with DADA2 parameters *maxN=0, truncQ=2, rm*.*phix=TRUE and maxEE=2*. The *assignTaxonomy* method of DADA2 was used to assign a genus-level taxonomic classification to each of the ASVs using the Silva v. 138 database (20) as a reference.

ASV count tables were rarefied to a depth of 10,000 reads per sample, and samples with less than 10,000 reads were not considered in the analysis. The rarefied count matrices were used to calculate the Chao1 and Shannon diversity indexes at the ASV level for each sample using the alpha_diversity module in the sci-kit bio (v. 0.5.6) python library.

### Statistical analysis

ANOSIM and PERMDISP analyses between sample groups were done using the sci-kit bio (v. 0.5.6) python library based on the Bray-Curtis (BC) dissimilarity matrices of the ASV relative abundances per sample. Principal component analysis of the BC dissimilarity matrix was carried out using the sci-kit learn (v. 0.21.3) python library.

Differential abundances of ASVs according to liver abscess diagnosis were determined using the generalized additive model for location, scale and shape (GAMLSS) implemented in metamicrobiomeR with a zero-inflated beta distribution (21). For each study, data from all timepoints and treatments were considered simultaneously, with animal id included as a random variable for longitudinal analysis and adjusting for treatment group as a fixed effect. Only ASVs with a mean relative abundance across samples higher than 10^−4^ and prevalence in at least 5% of samples were tested. The 10^−4^ threshold was selected given the higher dominance of technical noise for low abundance taxa in comparable microbial communities (22). A false discovery rate (FDR) threshold of 0.05 was used to filter ASVs associated with abscess state. Fisher’s exact tests were used to determine significantly enriched or depleted bacterial families among differentially abundant ASVs.

### Data availability

Raw reads were submitted to the Sequence Read Archive with accession PRJNA686084.

## Results

Out of 200 steers assigned to the S1-control treatment in study 1, 68% showed abscesses at the time of necropsy. This was significantly higher than the 29% abscessed animals in the S1-tylosin group (Fisher’s exact test p-value = 5×10^−15^). In study 2, 43% of animals in the S2-control group developed abscesses compared to 35% in the S2-narasin group (p-value = 0.12). Fecal samples were collected from normal and abscessed steers from the control and antibiotic treatments to investigate 1) whether there are differences in fecal microbiome composition according to liver abscess state, 3) whether those differences are more prevalent at early versus late stages of feeding, and 3) whether any differences are modulated antibiotic treatment.

Out of the 21 sampled animals from the S1-control treatment, 15 were abscessed and 6 were normal at the time of necropsy. Out of the 22 sampled animals from the S1-tylosin treatment, 8 were abscessed and 14 had normal livers. For both the S2-control and S2-narasin treatment groups, samples were collected for 15 normal and 5 abscessed animals. For each sampled animal, samples were collected at four (study 1) or six (study 2) timepoints during the study as indicated by the red markers in Figure 1.

**Figure 1.**
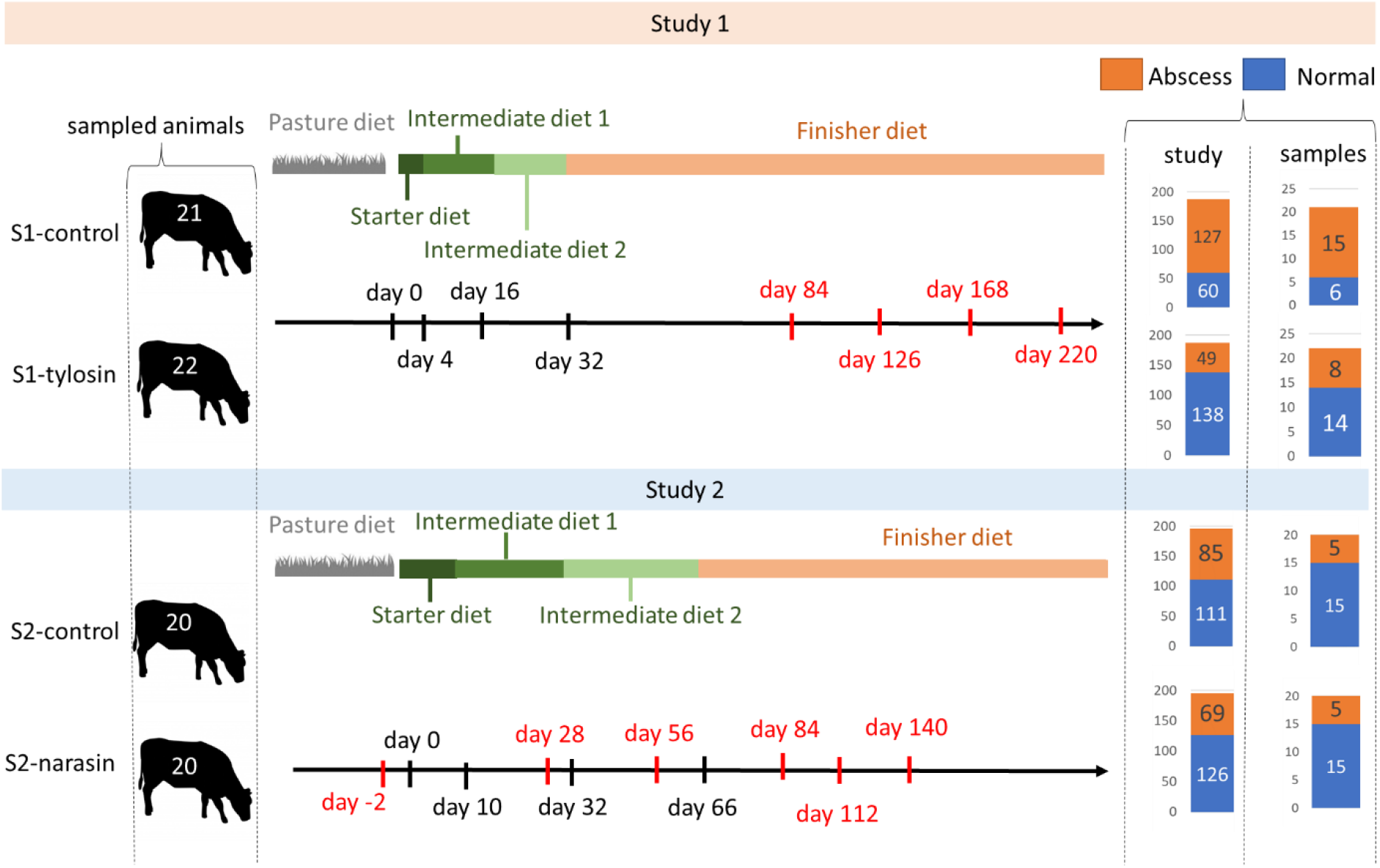
Treatments and collected samples for study 1 and study 2. Overview of diet schedules, treatments, and numbers of sampled animals in two clinical studies. The black arrows represent time. Black markers indicate the days in which diets were changed; red markers show days fecal samples were collected. The bar plots on the right show, for each treatment in each study, the number of animals diagnosed with liver abscesses at the time of necropsy, as well as the number of animals from which fecal samples were collected according to their liver abscess status.

After processing of the raw reads, ASV counts for 162 fecal samples from study 1 and 354 fecal samples from study 2 were analyzed. At the phylum level (Figure 2A, B), fecal microbial communities were dominated by Firmicutes (61 and 51 % for studies 1 and 2, respectively) followed by Bacteroidetes (29 and 36%), Proteobacteria (6 and 7%) and Spirochaetes (1% in both studies). Regardless of abscess state at the time of necropsy, ASV alpha-diversity quantified by both the Chao1 and Shannon’s indexes showed a trend of increasing diversity as a function of time in study 1 (Figure 2C, D). In study 2 a similar pattern was observed during the periods that the animals were being fed either the intermediate diets (days 28 to 56), or the finisher diet (days 84 to 140), regardless of abscess state. Nevertheless, substantial drops in diversity were observed consistent with the major diet transitions in the study (Figure 2E, F). In study 2, the drop in diversity from day -2 to day 28 was accompanied by a slight decrease in the abundance of Firmicutes and an increase in Proteobacteria and Spirochaetes. On the other hand, the decrease in diversity following the transition to the finisher diet was accompanied by an increase in Bacteroidetes (Figure 2B). Based on these results, we conclude that the alpha diversity of the fecal microbiome of steers in the feedlot decreased upon diet changes but increased the longer the animals remained on a given diet.

**Figure 2.**
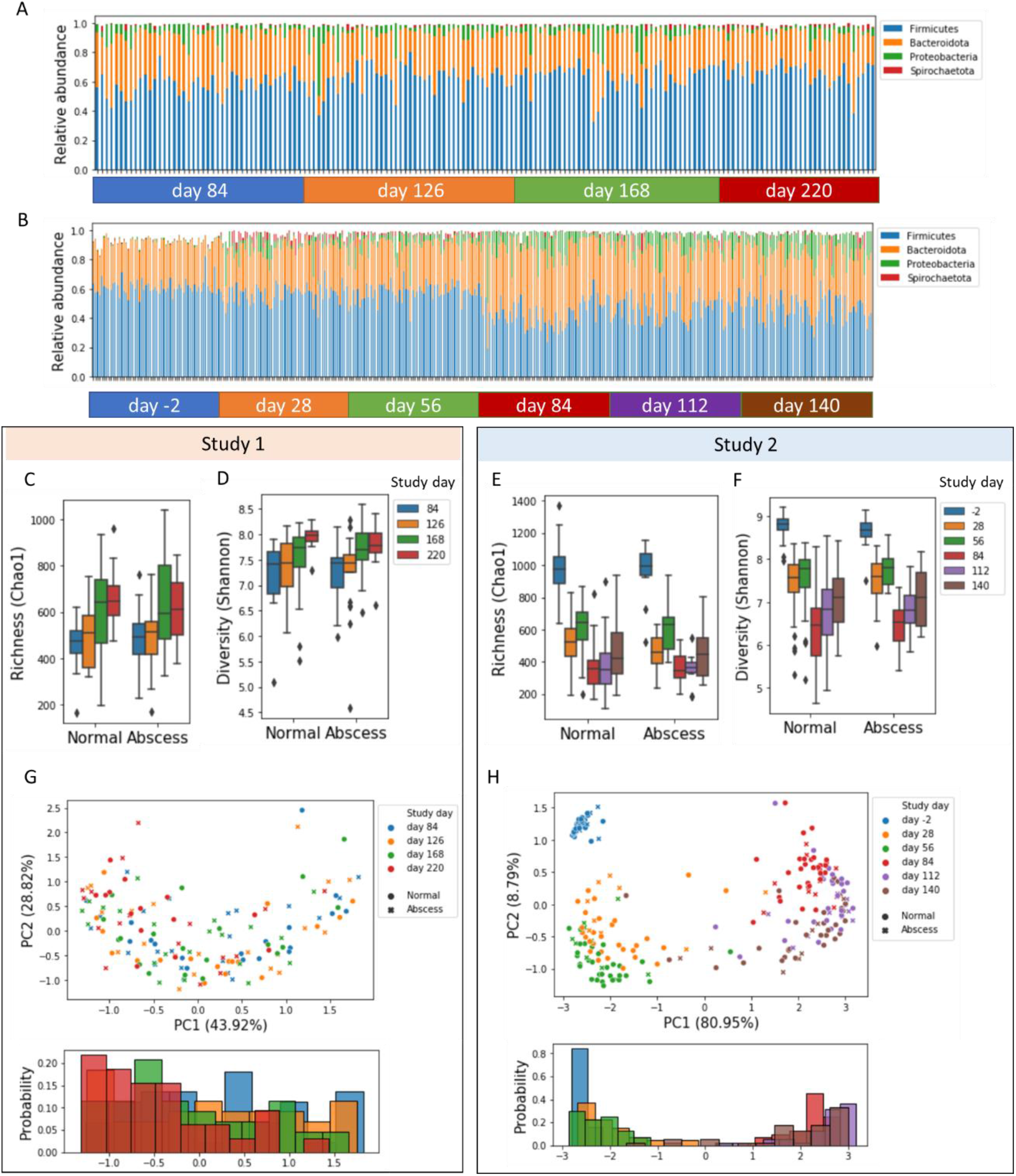
Composition of the fecal microbiome of steers in study 1 and study 2. **A**. Relative abundances of major phyla of bacteria detected in samples passing quality thresholds in study 1. Each bar represents an individual sample. Samples are grouped according through the time of sample collection. **B**. Like A but for study 2. **C, D**. the Chao1 and Shannon alpha diversity indexes for ASVs in samples from normal or abscessed steers at different study days from study 1. Boxes represent the interquartile range and whiskers extend to minimum and maximum values, except for points outside 1.5 times the interquartile range. **E, F**. Like C, D but for study 2. **G**. The two first principal components (PCs) for the Bray-Curtis dissimilarity matrix of the relative abundances of ASVs in samples from study 1. Colors represent different fecal sample collection times and symbols indicate abscess diagnosis of corresponding animals. Numbers in parentheses indicate the variance explained by the corresponding PC. The histogram shows the distribution of samples from different times on the first PC. **H**. Like G but for study 2.

Principal component analysis of the Bray-Curtis dissimilarities between relative ASV abundances across samples showed that, in both studies, the fecal communities varied primarily as a function of time on diet (Figure 2G, H). These results were supported by significant p-values (<0.05) for ANOSIM analyses comparing the fecal communities between samples collected at different timepoints (Supplementary Table 1). For samples collected at individual timepoints, ANOSIM between samples from the S1-control and S1-tylosin treatments from study 1 did not detect significant differences in community composition. In study 2, however, we observed significant differences between the S2-control and S2-narasin treatments at all timepoints except for day -2, which occurred before treatments were applied (Supplementary Table 2).

Taken together, regardless of the different diet regimes and sampling times between both studies, the fecal microbiomes showed similar patterns in terms of the abundances of major phyla and changes in alpha and beta diversity; with a dominant role of diet shifts and time on diet, and a detectable effect of narasin treatment but not tylosin treatment on fecal microbiome composition.

To investigate the potential association between fecal microbiome composition and liver abscess diagnosis at the time of necropsy, we carried out ANOSIM analyses at each timepoint between samples from abscessed and normal animals. The results revealed no significant differences in the fecal microbiome for any of the timepoints for the control animals of both studies (Figure 3A, B). In study 1, the microbiome dissimilarity (measured by the ANOSIM R score) between abscessed and normal cattle increased between day 84 and day 220. For study 2, the dissimilarity tended to decrease from day -2 to day 140. These patterns were different in the antibiotic treatments from both studies as follows. For tylosin treated animals in study 1 the largest dissimilarity between normal and abscessed steers was observed on day 128 (Figure 3C, p-value=0.02). For narasin treated animals from study 2 the largest difference was observed on study day 28 (Figure 3D, p-value=0.003) and, similar to study 2 controls, dissimilarity tended to decrease through the length of the study. Interestingly, in both the S2-control and S2-narasin treatments there were relatively large (although not statistically significant—p-value>0.05) differences between abscessed and normal steers on day -2 of the study; that is, before the first intermediate diet with or without antibiotic was introduced. Thus, it is possible that differences between animals prior to the start of the experiment might be associated with liver abscess susceptibility. Collectively, these observations suggest an interaction between time on diet, antibiotic treatment, and abscess diagnosis on the fecal microbiome composition of steers in the feedlot.

**Figure 3.**
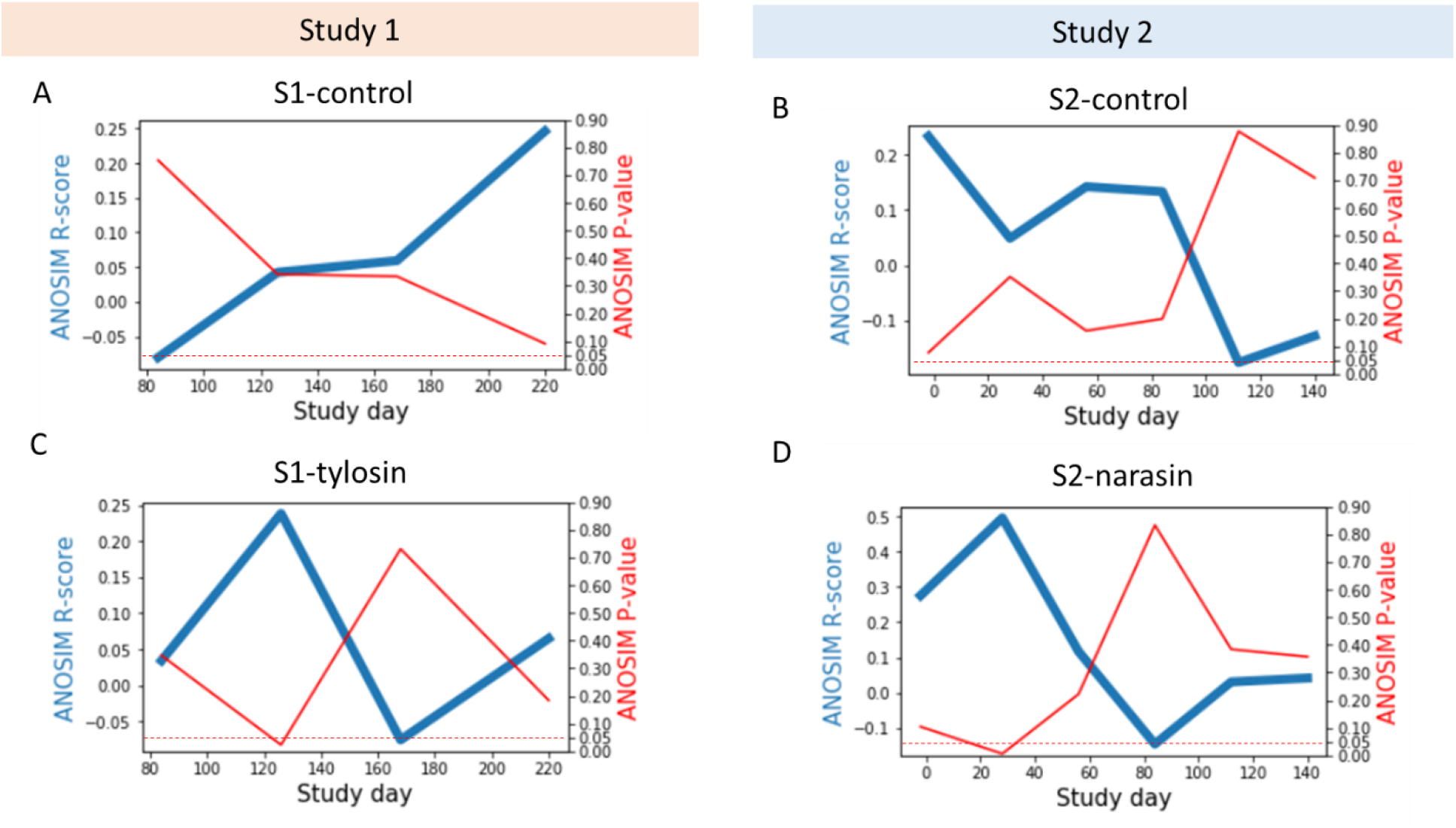
Similarities between the fecal microbiomes of abscessed and normal steers. **A**. the ANOSIM R scores (blue line, left y-axis) for the comparison between samples from normal and abscessed animals from the S1-control treatment at different timepoints. The red line and right y-axis show the corresponding p-values. The dashed line indicates an ANOSIM p-value of 0.05. **B-D**. Like A but for animals from the S2-control, S1-tylosin and S2-narasin treatments.

To better understand the effect of time on diet, treatment, and liver abscess on fecal microbial communities, we compared dispersion between microbiome profiles for samples from the same treatment between animals with and without abscesses at different timepoints throughout both studies. To do so, we plotted the average Bray-Curtis dissimilarity between pairs of samples in each of the groups as a function of time (Figure 4). In the control treatments of both studies, we saw little difference in the pairwise dissimilarity of abscessed and normal steers (Figure 4A, B), with dispersion between samples remaining similar through the studies. In contrast, for antibiotic treated animals, we observed a clear separation, with abscessed animals having a higher dispersion than normal steers on day 128 for study 1 (Figure 4C, PERMDISP p-value=0.036) and on day 28 for study 2 (Figure 4d, PERMDISP p-value=0.01). The results indicate contrasting patterns of microbiome variability between steers that developed or not liver abscesses while receiving preventative antibiotics. The lower variability observed for normal animals receiving antibiotics also tended to be lower than that observed for normal animals in the control treatments at the corresponding timepoints (p-value=0.098 and 0.044 for study 1 and 2 respectively). Thus, the detected differences in microbial community structure between abscessed and normal animals in the s1-tylosin and s2-narasin treatment, are largely a result of differences in microbiome variability. The results suggest that effective prevention of liver abscesses by antibiotics might be accompanied by a more uniform composition of the fecal microbiomes of steers at specific timepoints in the feedlot.

**Figure 4.**
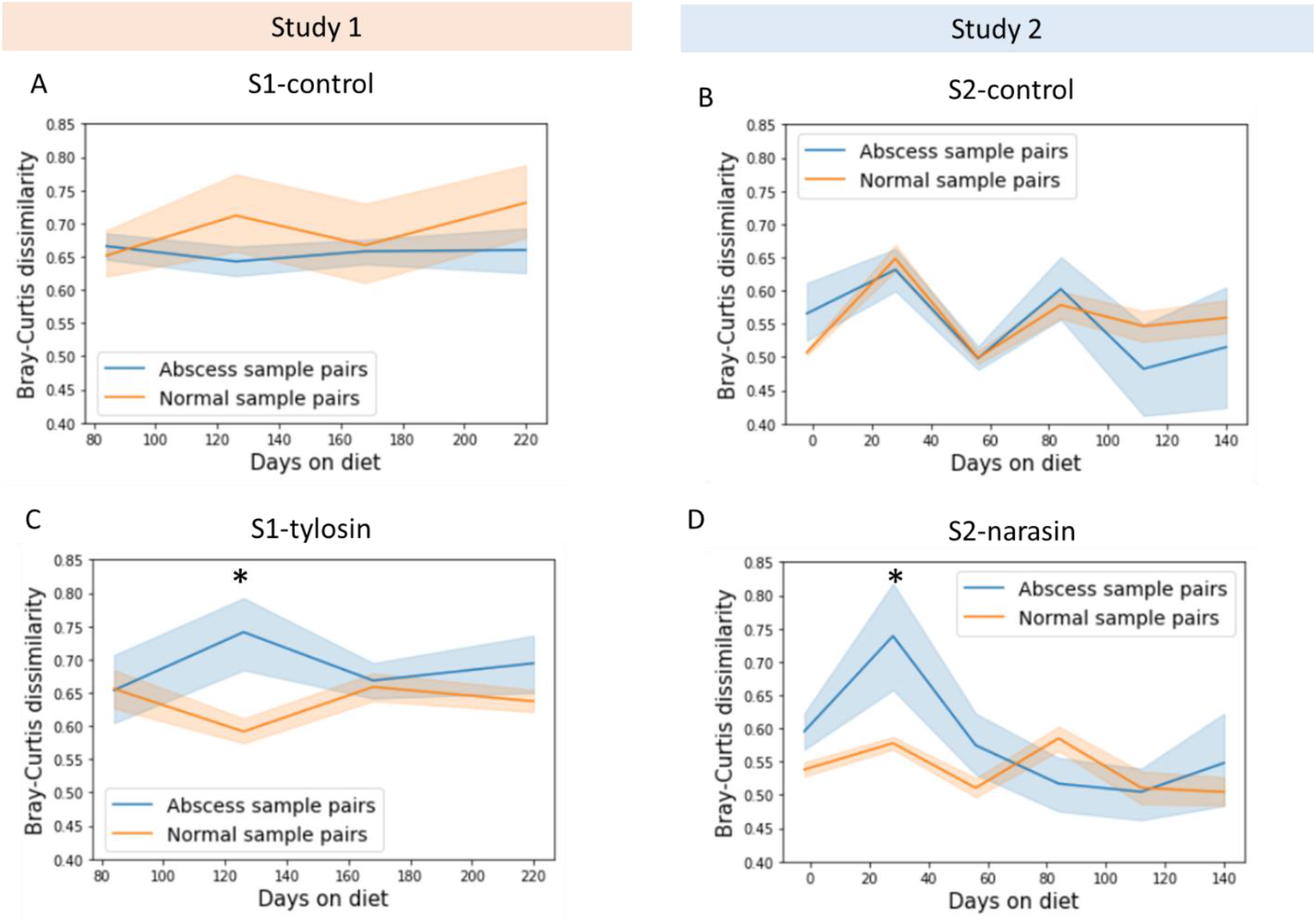
Dispersion of beta-diversity values of abscessed and normal animals. **A**. The average Bray-Curtis dissimilarity between all pairwise comparisons between samples from the S1-control treatment as a function of time on diet. Colors indicate samples from abscessed or normal steers. The bands around each line indicate the 95% confidence interval for the mean. **B-D**. Like A but for samples from the S2-control, S1-tylosin and S2-narasin treatments. * in panels C and D indicate a p-value < 0.05 for PERMDISP tests between abscess and normal beta-diversity values at the indicated timepoints.

Finally, we investigated microorganisms whose relative abundances were higher in animals that developed abscesses and those that did not, in both studies. For this, we applied the GAMLSS model implemented in metamicrobiomeR (21). After adjusting for the effect of treatment and considering that repeated measures from the same animals were done in the longitudinal design, we obtained 35 and 44 ASVs negatively associated with abscesses in studies 1 and 2 respectively (FDR<0.05), and 41 and 56 ASVs positively associated with abscesses (Supplementary tables 3, 4). Notably, while 663 out of 1424 ASVs tested for differential abundance were shared between both studies, none of the significant ASVs were found in common between the studies. Looking at the taxonomic classification of differentially abundant ASVs from both studies showed similar patterns for the number of ASVs from different bacterial families associated with normal or abscessed animals (Figure 5A, B). This likely reflects similarities in overall fecal microbiome composition between studies, as only a handful of families were significantly enriched with ASVs associated with liver abscess (Fisher’s exact test p-value < 0.05, arrows on Figure 5). In other words, in most cases families with more ASVs also had more abscess-associated ASVs. Thus, while the individual studies pointed to specific microbes associated with liver abscesses, these patterns were not shared between studies.

**Figure 5.**
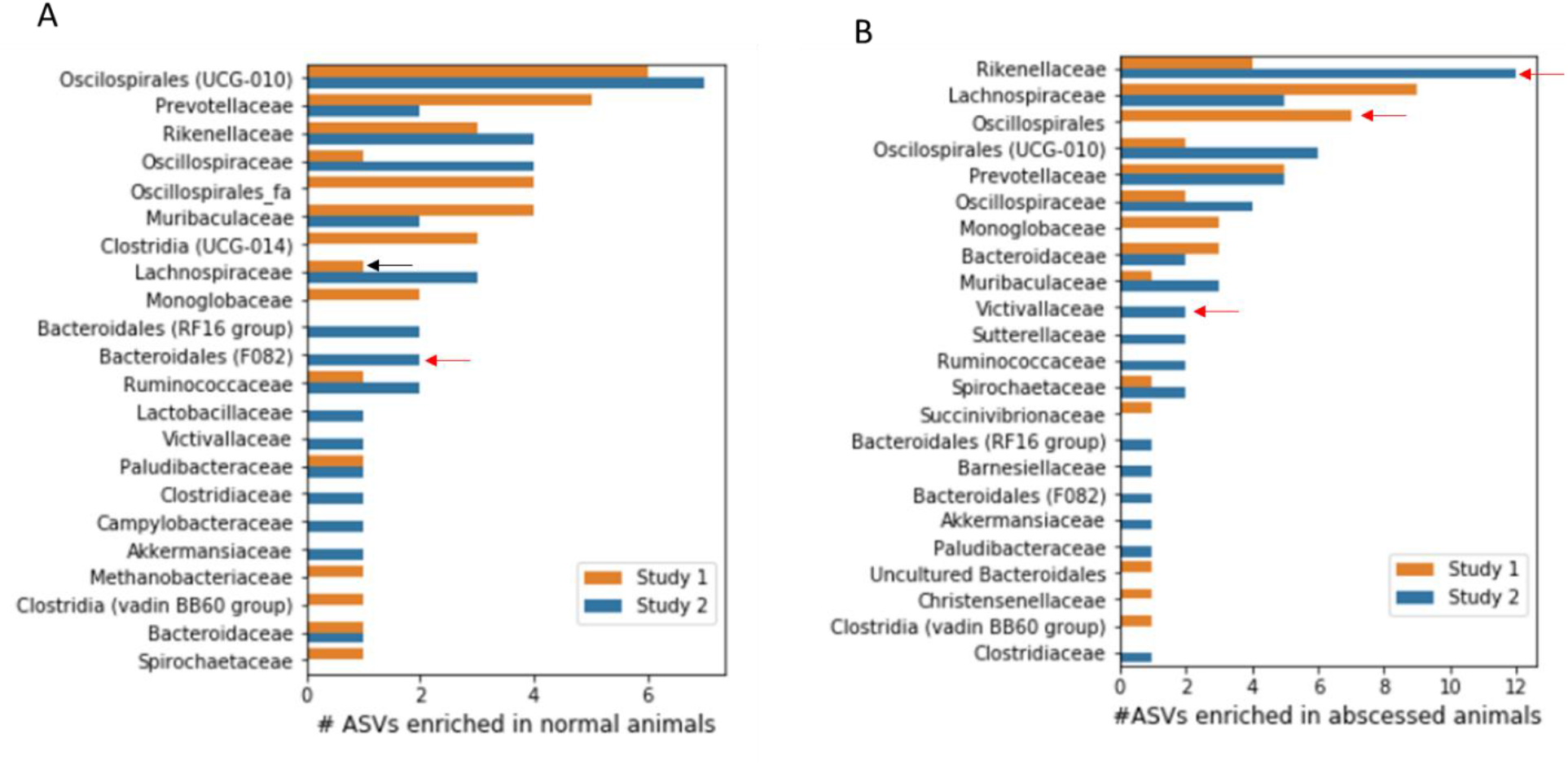
Taxonomic classification of ASVs associated with liver abscesses. **A**. Number of ASVs per bacterial family with higher relative abundances in normal animals than abscessed animals. **B**. Number of ASVs per bacterial family with higher relative abundances in abscessed animals than normal animals. In both panels colors indicate results for two different studies. Red arrows indicate families with a significant enrichment of differentially abundant ASVs (Fisher’s exact test p-value < 0.05). Black arrows indicate families significantly underrepresented among differentially abundant ASVs.

## Discussion

In this manuscript, we describe the composition of the fecal microbiome of feedlot steers that developed or not liver abscesses, fed with or without antibiotics, using longitudinal data from two clinical studies. Given the easier access to fecal samples compared to blood or rumen contents, the identification of fecal biomarkers for liver abscess risk would facilitate managing of the disease in the field. For instance, to only treat animals at a higher risk of developing abscesses or inform cases in which preventative antibiotics have not been effective. Our results showed subtle but significant differences in the fecal microbiomes of animals with and without abscesses. These included differences in community structure and variance. The observed differences were more marked towards early timepoints following the transition from pasture to a high energy diet in the feedlot. Interestingly, samples collected 2 days before the start of study 2 showed larger differences between animals that developed and did not develop liver abscesses than samples collected at late timepoints on a high concentrate diet. This suggests that fecal microbiomes might differentiate animals more susceptible to liver abscesses even before entering the feedlot. Our analysis also revealed larger microbiome differences between abscessed and normal steers when these had been treated with tylosin (study 1) or narasin (study 2).

The observed differences coincided with timepoints for which samples from animals that developed abscesses were more different from each other than those from normal animals in the antibiotic treatments. The observed patterns suggest that tylosin and narasin led to a reduced dispersion of microbial communities when they were effective at preventing abscesses. Stochastic changes in microbial community composition have been recently discussed in the context of type 1 diabetes (23), obesity (24), alcoholism (25), smoking (26), and liver cirrhosis (27) among others, illustrating how animal microbiomes may exhibit increased dispersion under stress (28). We observed strong directional changes in the abundance of major taxonomic groups as a function of diet changes and time on diet. These changes were characterized by a drop in microbial diversity upon switching diets, followed by a steady increase while the animals were kept on the same diet. These transition periods are known stressors for cattle entering the feedlot (29) and given the observed early microbiome differences of animals that developed or not abscesses reported here, it is likely that managing that initial stress contributes to liver abscess prevention. Indeed, a lower overall prevalence of abscesses among animals in the control treatment was seen in study 2, for which the diet transition period was twice longer than in study 1.

Even though a substantial number of ASVs from diverse families were found to be associated with liver abscess diagnosis at the time of necropsy in each of the studies, we found no overlap between both studies. This is likely related to the differences between treatments and diets used in both studies. Thus, while we saw some similarities in the nature of beta diversity differences associated with abscesses between studies, the abundance of individual microorganisms may be limited in its capacity to robustly predict liver abscesses across different feedlots. Given the number of independent samples analyzed in this study (∼20 animals per treatment), building and testing microbiome-based predictive models is beyond the scope of this study. It is possible that a focus on functional content, gene activity or metabolite levels shows more robust associations with abscesses than ASV abundances, or that a larger number of individuals needs to be sampled to discover robust associations (30). Nevertheless, our results point to early differences in the hindgut environment of animals that develop liver abscesses, leading to diverging fecal microbial communities.

